# Developmental Effects of Lithium and Zinc in Planaria

**DOI:** 10.1101/2024.08.30.610554

**Authors:** Alberto Molano

## Abstract

An investigation was carried out of the stage-specific effects of lithium and zinc on planaria embryologic development. These two reagents had dramatically different effects on the morphological development of planaria: Zinc induced the formation of bifurcations or trifurcations of the head region. In contrast, lithium induced the formation of round masses of cells. The earliest embryological stages (S1 and S2, cryptic larva formation) exhibited extreme sensitivity to lithium, since the lethal dose was at least one order of magnitude lower than for subsequent stages 3-5, suggesting that beta-catenin 1 expression is tightly regulated during the initial embryologic stages and that even slightly increased activity is lethal.

## Introduction

Zinc sulphate and lithium chloride have been traditionally used in developmental biology to modify embryonic cell fates. Lithium chloride is a classical “vegetalizing” (posteriorizing) agent, while zinc sulfate “animalizes” (anteriorizes) embryos. These reagents perturb the development of a multitude of organisms. In invertebrates like sea urchins, lithium acts as a posteriorizing agent, expanding the endomesoderm and eliminating the apical organ region, while zinc acts as an anteriorizing agent, generating embryos with no or reduced endomesoderm (1, 2). In vertebrates like *Xenopus laevis*, the effects of lithium on embryonic development depend on the dose and stage of administration (3, 4).

Biochemical experiments have demonstrated that the molecular target of lithium is glycogen synthase kinase 3 beta (GSK-3β), a key component of the Wnt signaling pathway (5). Lithium inhibition of GSK-3β (*K*_*i*_ = 2.1 mM) prevents beta catenin degradation by the destruction complex and permits its accumulation in the nuclei, where it binds to the TCF/LEF transcription factor to modulate gene expression. The molecular target(s) of zinc is (are) unknown.

These initial events trigger complex downstream responses involving other transcription factors that are sequentially activated to specify cell fates at defined moments of embryonic development. Such developmental gene regulatory networks (dGRNs) have been particularly well studied in sea urchins like *Strongylocentrotus purpuratus* (2). In this species, lithium is a key activator of the endomesodermal gene regulatory network, leading to excessive endomesoderm and elimination of apical pole genes. Sea urchin embryos treated with lithium upregulate endomesoderm-specific genes like *Brachyury, gata-e, foxa, hox11/13b, notch, wnt8, krl*, and *endo16*, while strongly downregulating apical ectodermal genes like secreted frizzled protein 1/5. The result is an amorphous mass of endoderm and mesoderm. In contrast, zinc sulphate blocks endomesoderm specification while upregulating apical ectodermal and neural markers. The result is an embryo that is arrested as a hollow ball of ectodermal cells (2, 6).

No reports exist of the morphological effects of lithium and zinc on planaria embryos. Here, an investigation was conducted of the stage-specific effects of lithium and zinc on planaria embryologic development.

## Materials & Methods

### Planaria

Planaria were collected from Río Sáchica, a stream in Villa de Leyva, Boyacá, Colombia (5.6365° N, 73.5271° W; altitude 2,149 meters above sea level) (7). Morphologically they appear to be *Girardia sp*. Worms reproduced sexually and were maintained in a tank filled with bottled stream water and oxygenated with an aquarium pump. They were fed raw beef liver once a week.

### Eggs

The moment of egg deposition was determined visually by means of frequent inspections and by the color of the capsule (amber immediately after deposition but turning black after a few hours).

### Lithium chloride and Zinc Sulphate

Lithium chloride anhydrous (99.06% purity, lot # B463812306) and zinc sulphate heptahydrate (99.82% purity, lot # B382802109) were purchased from Loba Chemie (Mumbai, India).

### Treatments

The planaria embryos studied in this paper take approximately 20 days to develop from deposition to hatching (8). To investigate the stage-specific effects of lithium and zinc during development, lithium chloride and zinc sulphate were added during three intervals: (A) Days 1-4, (B) 5-8, and (C) 9-12. By day 13 the basic planaria body plan has already been established (8), so no experiments were conducted beyond day 12. Freshly prepared solutions of the reagents were added to the eggs (n=5) at the specified time points, followed by extensive washing with water. Images were captured with a smartphone and a Swift 380T microscope (Xiamen, China).

## Results

Planaria embryologic development is usually divided into 8 stages (8). Some critical morphological and gene expression events have been identified in other planaria species like *Schmidtea polychroa* and were used as the basis for this study (8). The planaria embryos studied in this paper hatch 20 days after deposition on average. To match developmental events with days post-deposition, eggs were punctured at various time points (8) (Figure 1).

**Figure 1.**
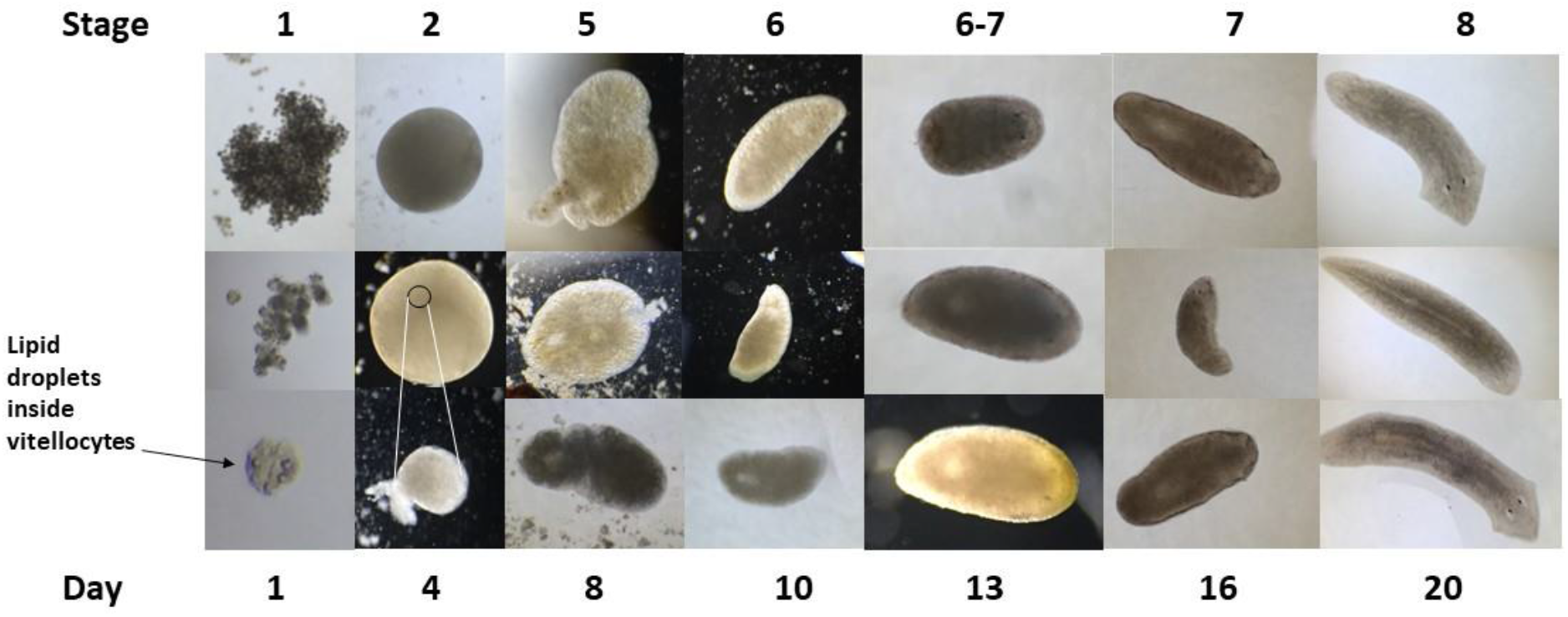
Embryologic stages. Eggs were gently punctured and dissected at the indicated time points/stages to analyze embryo morphology. Notice the lipid droplets inside vitellocytes in stage 1 embryos. The upper two photographs of stage 2 embryos are fixed specimens (4% formaldehyde overnight) showing the entire content inside the egg capsule, whereas the lower picture shows an embryo dissected in an unfixed specimen. The small circle in the middle picture indicates the approximate size of the embryo in relation to the egg.

To investigate the stage-specific effects of lithium and zinc during development, lithium chloride and zinc sulphate were added during three intervals: (A) Days 1-4, (B) 5-8, and 9-12. By day 13 the basic planaria body plan has already been established (8), so no experiments were conducted beyond day 12.

Based on published results with other species like sea urchins, 50- and 25-mM lithium chloride and 0.5- and 0.1-mM zinc sulphate were tested initially (1, 2). However, these lithium concentrations were 100% lethal, while the zinc concentrations had no effect whatsoever. Lithium chloride was then titrated down (10, 5, 2.5, and 1 mM) and the optimal teratogenic concentration was found to be 5 mM, which resulted in approximately 20% of worms with malformations when added on days 5-8. Remarkably, however, this lithium concentration continued to be 100% lethal when added on days 1-4. Hence, a second lithium down titration was undertaken (1, 0.5, 0.4, 0.35, 0.3, and 0.25 mM) specifically for days 1-4. These experiments showed that 1- and 0.5-mM lithium were 100% lethal on days 1-4, whereas 0.35, 0.3, and 0.25 mM had no effect. The optimal teratogenic concentration on days 1-4 was determined to be 0.4 mM. A striking difference was noticed in the susceptibility to lithium of the earliest (S1 and S2) embryonic stages when compared to subsequent ones (S3-S5), since the lethal dose for days 5-8 (stages S3-S5) was 10 mM, whereas the lethal dose on days 1-4 (stages S1 and S2) was only 0.5 mM, which is 20-fold lower.

In terms of developmental abnormalities, lithium induced the formation of round masses of cells when embryos were exposed on days 1-4 and 5-8 (Figure 2). Exposure on days 9-12 produced no effects (Figure 2). In contrast, zinc induced characteristic bifurcations or trifurcations of the head region when embryos were exposed in days 1-4 and 5-8. On days 9-12 there were no effects (Figure 2).

**Figure 2.**
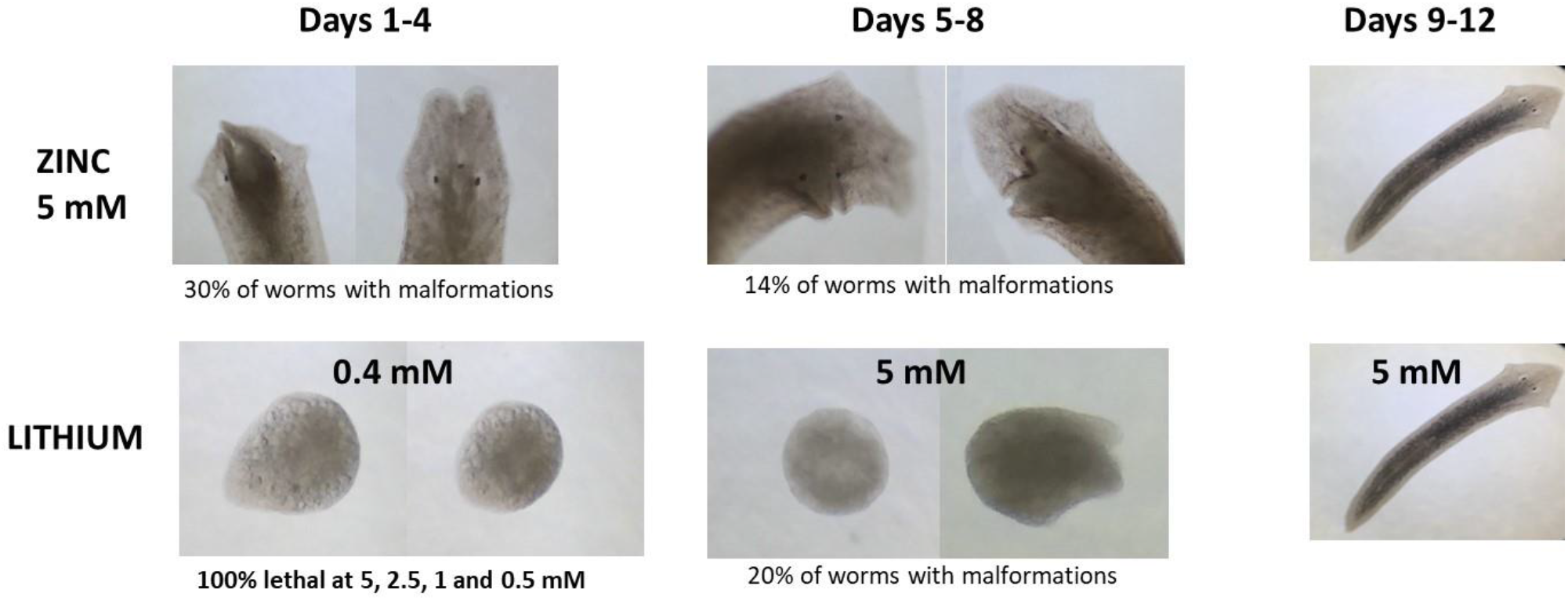
Developmental abnormalities induced by lithium and zinc in planaria. Lithium induced the formation of round masses of cells when embryos were exposed on days 1-4 and 5-8 (Figure 2). Exposure on days 9-12 produced no effects (Figure 2). In contrast, zinc induced characteristic bifurcations or trifurcations of the head region when embryos were exposed in days 1-4 and 5-8. On days 9-12 there were no effects. A striking difference was noticed in the susceptibility to lithium of the earliest (S1 and S2) embryonic stages when compared to subsequent ones (S3-S5), since the lethal dose for days 5-8 (stages S3-S5) was 10 mM, whereas the lethal dose on days 1-4 (stages S1 and S2) was only 0.5 mM, which is 20-fold lower.

## Discussion

Zinc sulphate and lithium chloride have been traditionally used in developmental biology to modify embryonic cell fates. Lithium chloride is a classical “vegetalizing” (posteriorizing) agent, while zinc sulfate “animalizes” (anteriorizes) embryos. Lithium inhibits glycogen synthase kinase 3 beta (GSK-3β), prevents beta catenin degradation by the destruction complex, leading to its accumulation in the nuclei, where it binds to the TCF/LEF transcription factor and, in species like the sea urchin *Strongylocentrotus purpuratus*, activates the endomesodermal gene regulatory network, leading to excessive endomesoderm and elimination of apical pole genes (2, 5). In contrast, zinc sulphate blocks endomesoderm specification while upregulating apical ectodermal and neural markers. In the amphibian *Xenopus*, LiCl-treated embryos highly upregulate early dorsal fate determinants expressed in the Spemann organizer like the homeobox genes siamois 1 (sia1) and goosecoid (gsc); BMP antagonists noggin (nog) and chordin (chrd); canonical Wnt regulators frizzled 8 (fzd8) and loc3888630 (tiki1); and the secreted tyrosine kinase pkdcc.1 (4).

No reports exist of the morphological effects of lithium and zinc on planaria embryos. Here, an investigation was conducted of the stage-specific effects of lithium and zinc on planaria embryologic development. Results showed that lithium induced the formation of round masses of cells when embryos were exposed on days 1-4 and 5-8, suggesting activation of the endomesodermal gene regulatory network, excessive endomesoderm formation, and downregulation of ectodermal genes. In contrast, zinc induced characteristic bifurcations or trifurcations of the head region when embryos were exposed on days 1-4 and 5-8, suggesting upregulation of anterior polarity genes (Figure 2). These morphological anomalies agree with the known anteriorizing and posteriorizing effects observed in other species.

